# Regional Centromere Configuration in the Fungal Pathogens of *Pneumocystis* Genus

**DOI:** 10.1101/2023.05.12.540427

**Authors:** Ousmane H. Cissé, Shelly Curran, H. Diego Folco, Yueqin Liu, Lisa Bishop, Honghui Wang, Elizabeth R. Fischer, A Sally Davis, Christian Combs, Sabrina Thapar, John P. Dekker, Shiv Grewal, Melanie Cushion, Liang Ma, Joseph A. Kovacs

## Abstract

Centromeres are constricted chromosomal regions that are essential <u>for</u> cell division. In eukaryotes, centromeres display a remarkable architectural and genetic diversity. The basis of centromere accelerated evolution remains elusive. Here we focused on *Pneumocystis* species, a group of Mammalian-specific fungal pathogens that form a sister taxon with that of the *Schizosaccharomyces pombe*, an important genetic model for centromere biology research. Methods allowing reliable continuous culture of *Pneumocystis* species do not currently exist, precluding genetic manipulation. CENP-A, a variant of histone H3, is the epigenetic marker that defines centromeres in most eukaryotes. Using heterologous complementation, we show that the *Pneumocystis* CENP-A ortholog is functionally equivalent to CENP-A^Cnp1^ of *Schizosaccharomyces pombe*. Using organisms from a short-term *in vitro* culture or infected animal models and ChIP-Seq, we identified CENP-A bound regions in two *Pneumocystis* species that diverged ∼35 million years ago. Each species has a unique short regional centromere (< 10kb) flanked by heterochromatin in 16-17 monocentric chromosomes. They span active genes and lack conserved DNA sequence motifs and repeats. These features suggest an epigenetic specification of centromere function. Analysis of centromeric DNA across multiple *Pneumocystis* species suggest a vertical transmission at least 100 million years <u>ago</u>. Common ancestry of *Pneumocystis* and *S. pombe* centromeres is untraceable at the DNA level but the overall architectural similarity could be the result of functional constraint for successful chromosomal segregation.

**Significance Statement:** *Pneumocystis* species offer a suitable genetic system to study centromere evolution in pathogens because of their phylogenetic proximity with the nonpathogenic yeast *Schizosaccharomyces pombe*, a popular model for cell biology. We used this system to explore how centromeres have evolved after divergence of the two clades ∼460 million years ago. To address this question, we established a protocol combining short-term culture and ChIP-Seq to characterize centromeres in multiple *Pneumocystis* species. We show that *Pneumocystis* have short epigenetic centromeres that function differently from those in *S. pombe*.

**One sentence summary:** Insights into the formation of genic centromeres in fungal pathogens.

## Introduction

Centromeres are genomic locations where spindles attach during chromosomal segregation. They are essential for cell division. Errors in the chromosomal segregation can result in aneuploidy and cell division defects with disastrous consequences such as cancer. In fungi, centromeres are fragile sites that are often involved in karyotype variability (*1, 2*) and drug resistance (*3*). In most eukaryotes, centromeres are defined by the presence of CENP-A nucleosomes, a centromere specific histone H3 variant that is essential for the localization of known kinetochore components (*4*). There is a remarkable diversity in centromere structures ranging from sequence-dependent (point) centromeres to epigenetically-regulated (regional) centromeres.

Point centromeres are exemplified by *Saccharomyces cerevisiae* with genetically defined 125 bp long centromeres containing conserved DNA elements (I, II and III) that serve as anchors for the recruitment of the centromeric DNA binding factor 3 (CBF3) complex (*5, 6*). Regional centromeres are much longer, lack universally conserved DNA patterns, and are often made up of repetitive DNA (*7*). Regional centromeres are not strictly defined by DNA but are rather epigenetically regulated; however, which factors contribute to this epigenetic specification of centromeres remains elusive. In fungi, regional centromeres are classified as short (<20 kb, e.g. *Candida albicans* (*8*)), intermediate (40 – 110 kb; e.g. *Schizosaccharomyces pombe* (*9*) and long (150 – 300 kb, e.g., *Cryptococcus* (*10*)).

*Pneumocystis* is a genus of pathogenic fungi that exclusively infect mammals and remain unculturable. They belong to the Taphrinomycotina subphylum and form a monophyletic clade with fission yeasts *Schizosaccharomyces*, with a separation time estimated at ∼460 million years ago (*11, 12*). In addition to these two classes there are other classes, all of which are plant or soil adapted organisms. Compared to *S. pombe*, which is a tractable organism that is widely used as model for chromosome and cell biology*, Pneumocystis* species have streamlined genomes due to substantial gene losses during their transition to animal parasitism. Recently, the genomes of multiple *Pneumocystis* species have been sequenced (*12–15*). However, centromeres were not defined in these because they could not be predicted bioinformatically without reference for synteny based detection. Alternative methods such as chromosome conformation capture assay (4C), that have been used to predict centromere loci in fungal genomes (*16*) are not applicable to *Pneumocystis* because they require large quantities of pure DNA from synchronized cell cultures. No long-term culture system exists and during purification of *Pneumocystis* organisms from infected lungs it is virtually impossible to completely eliminate host cell DNA. *Pneumocystis* have retained CENP-A (*12, 15*), which presumably binds to centromeres, and most of the kinetochore proteins. Some data suggest that *Pneumocystis* centromeres differ structurally from those of *S. pombe*. For example, the heterochromatin protein Swi6/HP1 (*17*), which is required for <u>activation of</u> replication origins at the centromeres flanking regions (pericentromeres) has not been found in *Pneumocystis* genomes (*18*). The RNA interference (RNAi) pathway, which is essential to centromere function, has been lost in *Pneumocystis* (*19*), further supporting mechanistic difference with *S. pombe*.

In this work, we undertook to characterize *Pneumocystis* centromeres and compare them to *Schizosaccharomyces pombe* centromeres. <u>We found that *Pneumocystis* have small regional</u> <u>centromeres that are defined by CENP-A and flanked by heterochromatin, resembling those of *S. pombe*.</u> <u>However, *Pneumocystis* species lack orthologs for genes required for centromere function and</u> <u>maintenance in *S. pombe*, suggesting that its centromeres may function differently</u>.

## Results

### Survey of genes related to centromere function in Taphrinomycotina fungi

To understand the kinetochore evolution in Taphrinomycotina fungi, we screened the predicted proteomes of 14 species spanning six orders, *Pneumocystis* (*n* = 7), *Schizosaccharomyces* (*n* = 3), *Taphrina* (*n* = 1), *Saitoella* (*n* = 1), *Protomyces* (*n* = 1) and *Neolecta* (*n* = 1), as well as other fungi to have a broad overview of kinetochore evolution.

Using a combination of protein domain hidden Markov models and BLAST scans followed by phylogenetic assignment (see methods), we searched for homologs of CENP-A^Cnp1^, CENP-C^Cnp3^, CCAN (Constitutive Centromere Associated Network) and KMN (KNL-1/Mis12 complex/Ndc80 complex) proteins. We extended our searches to pathways required for centromere function and chromosomal segregation (heterochromatin, RNAi, and DNA methylation) (**Supplementary figure 1)**.

CENP-A, CENP-C, and outer kinetochore proteins are conserved across Taphrinomycotina. The CENP-A histones from *Pneumocystis* species, henceforth referring to as *pn*CENP-A, display a high overall conservation (89% average protein identity). Two genes of the inner kinetochore (CCAN) that are not found in *Pneumocystis* are innovations in *S. pombe* (*fta2* and *fta3* genes). The *fta2* and *fta3* gene products associate with the central core (cnt) and innermost repeats (imr) region of the centromere in *S. pombe* (*20*). No ortholog for the *mis*17 gene can be found in *Pneumocystis*. In *S. pombe*, *mis*17 encodes a member of the Mis6-Mal2-Sim4 multiprotein required for CENP-A recruitment (*21*). The *S. pombe* centromere-associated protein B genes (*cbp1*, *cbh1* and *cbh2*), which are derived from the domestication of *pogo*-like transposases (*22*) have no identified homologs in other fungi.

The chromosomal passenger complex (CPC) is a heterotetrametric complex composed of the Aurora B kinase Ark1 and three regulatory components Pic1 (inner centromere protein), Bir1p (Survivin) and Nbl1 (Borealin) which mediates chromosome segregation and cytokinesis (*23*). This pathway is conserved in *Pneumocystis* and *S. pombe*.

CENP-A recruiting complex, which includes Mis18 and Mis16 (*24*) as well as the CENP-A histone chaperone Scm3 required for CENP-A loading at the centromere (*25*) are conserved in *Pneumocystis*, as are the monopolin complex genes that coordinate the kinetochore microtubules attachment during mitosis (*26*).

The network for heterochromatin formation in *Pneumocystis* seems intact with the presence of the Clr4 methyltransferase complex (Clr4/SUV39H). *Pneumocystis* species have a single chromo domain containing protein that bear similarity with *S. pombe* Swi6 and Chp2 which are HP1 inparalogs (**Supplementary figure 2**).

DNA methyltransferases (DMTs) are divided into five classes (DNMT1, DNMT2, DNMT3/DRM, Masc1/RID and DNMT5)(*27*). Canonical DNA methylase (DMTs) are not detected in any of the seven sequenced *Pneumocystis* species (**Supplementary table 1**), which is consistent with previous studies (*28, 29*).

Overall, the *Pneumocystis* kinetochore (CCAN) protein catalog is relatively conserved and similar to those other Taphrinomycotina fungi.

### *pn*CENP-A^Cnp1^ localizes to the nucleus

Centromeres are identified by tracking CENP-A binding regions. Precise nuclear localization of CENP-A is required for accurate chromosomal segregation. To determine if *pn*CENP-A localizes in the nucleus, we used a anti – *Pneumocystis* CENP-A antibody, in conjunction with an anti- *Pneumocystis* antibody (*30*) and DAPI. Using indirect confocal immunofluorescence microscopy, we found that *pn*CENP-A displays a nuclear peripheral localization in replicating organisms from infected lung tissue and cultures (**Figures 1A and 1B**).

**Fig. 1.**
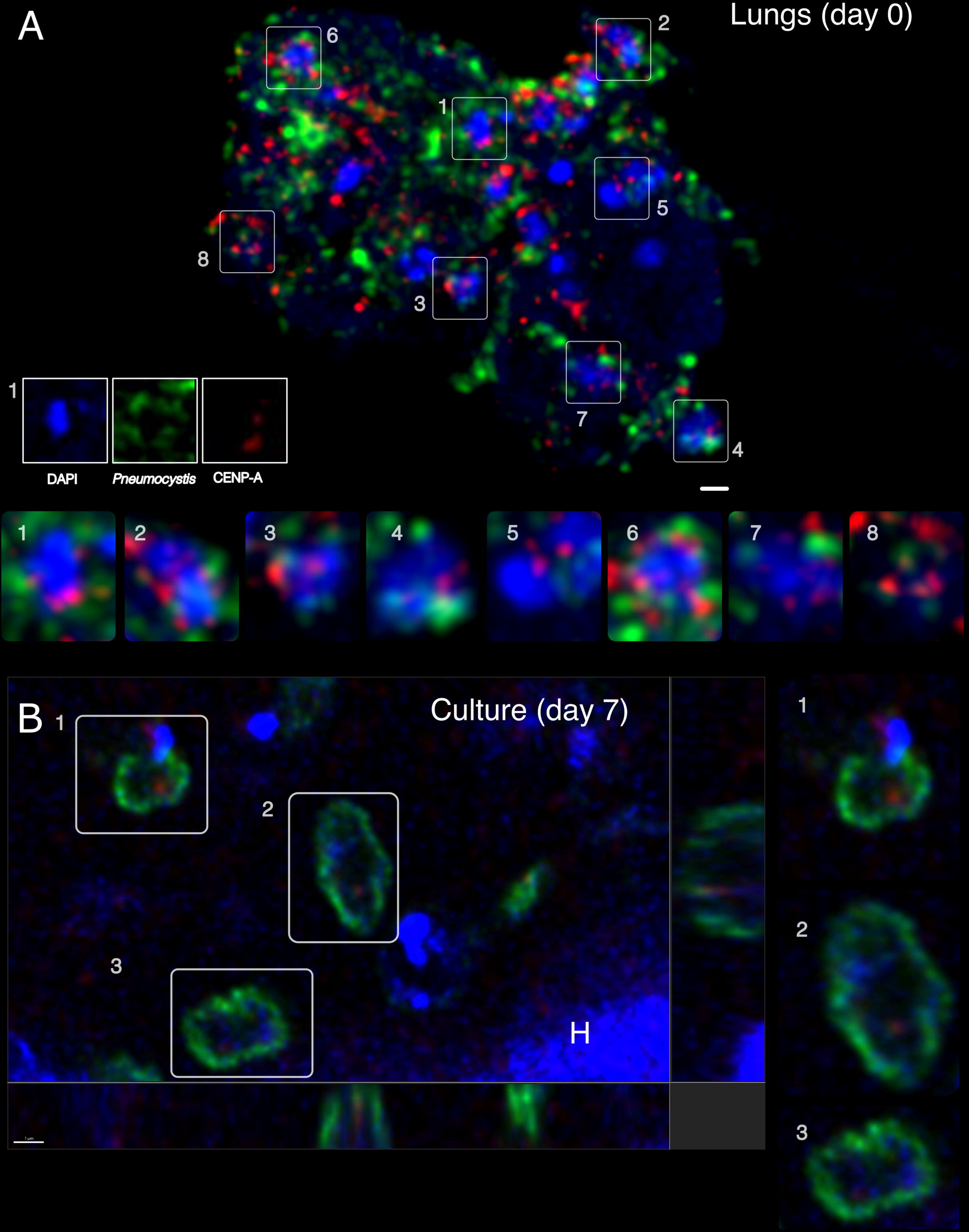
Nuclear peripheral localization of CENP-A in *Pneumocystis* cells. **(A**) confocal immunofluorescence microscopy shows that *pn*CENP-A (red) is present a nuclear periphery (blue) inside *Pneumocystis carinii* organisms (green). A cluster of organisms from an-infected rat lung tissue section is presented and labeled with an anti CENP-A antibody (red), an anti-*Pneumocystis carinii* antibody 7C4 (green) and 4′,6-diamidino-2-phenylindole (DAPI, blue). The image is a maximum projection of a 5 µm thick z-stack. Bottom panels 1-8 are magnified fields. Most anti *pn*CENP-A labels overlap or are close in proximity with DAPI (represented by a pink color produced by the overlap between red and blue colors). Some organisms are not labelled by 7C4 antibody (e.g., box 5), which suggest that the epitope is not expressed. This field does not contain any host cell (rat). Scale bar =L2 µm. (**B)** in plane and orthogonal views of immunofluorescence labelling of *P. carinii* organisms co-cultured with mammalian cells for seven days. Organisms were triple labeled using the same an anti CENP-A antibody (red), a different anti- *Pneumocystis* antibody (RAE7) targeting *Pneumocystis* major surface glycoproteins (green) and DAPI (blue). Panels 1-3 are magnified fields. CENP-A is located inside *Pneumocystis* cells at the periphery of the nucleus. Few c are only labeled with the DAPI and do not express detectable levels of CENP-A and major surface glycoprotein. A rat cell nucleus is labelled with the letter ‘H’. Scale bar =L1 µm.

### *pn*CENP-A is functional in *S. pombe,* and supports viability and centromere loading

To confirm that the putative *cenp-a* gene from *P. murina* (*PmCENP-A*) encodes a functional CENP-A protein, we expressed *PmCENP-A* in *S. pombe*. *P. carinii CENP-A* has only one amino acid difference with *P. murina* (**Figure 2A**). *PmCENP-A* was codon-optimized, expressed under the endogenous *S. pombe cnp1^+^*, the CENP-A ortholog, gene promoter and subcloned into a standard *LEU2*-based multicopy plasmid. In addition, the gene product was tagged with GFP at their N-terminus since it was shown that C-terminal tagging (*i.e.*, *cnp1^+^-GFP*) displays growth retardation (*31*). We first assessed the ability of *PmCENP-A* to rescue the lethality of *cnp1*Δ cells. We performed plasmid shuffling assays, in which *cnp1*Δ cells were initially kept viable by supplying a copy of *cnp1^+^* on a plasmid carrying the *ura4^+^* selection marker. A rescue plasmid carrying *PmCENP-A* and the *LEU2* selection marker was then transformed into the cells, followed by growth on medium containing 5-Fluoroorotic acid (FOA), which selects for loss of the *ura4^+^*-marked *cnp1^+^* plasmid (**Figure 2B**). As seen with control cells expressing the *S. pombe* GFP-CENP-A gene, we found that cells expressing *PmCENP-A* also fully support viability of *cnp1*Δ cells as they grew upon FOA selection (**Figure 2C**). We also observed no differences in growth between *cnp1*Δ cells expressing either *S. pombe* or *P. murina* CENP-A (**Supplementary figure 3**). We next asked if *PmCENP-A* can also rescue lethality of *cnp1-76* cells, a thermosensitive mutant, whose mutation T74M lies at the CATD (CENP-A targeting domain) region (*32*). As previously reported (*33*), we found that the *S. pombe* CENP-A tagged with GFP restored growth of *cnp1-76* cells at restrictive temperatures (i.e., 34°C). Importantly, *P. murina* counterpart was also able to rescue the thermosensitive phenotype (**Figure 2D**) proving the functionality of this gene. In interphase fission yeast displays the Rabl configuration, in which centromeres cluster to the spindle pole body while telomeres associate to each other near nuclear periphery (*34, 35*). As expected, expressing GFP-CENP-A from *S. pombe* resulted in cells displaying single fluorescence foci that colocalized with a *tetO* array inserted at *cen2* (NB: a TetR-tdTomato fusion is expressed to bind to the *tetO* array and this locus is referred to as *cen2-tetO-tdTomato*) (**Figure 2E**). Consistent with the mentioned rescue of *cnp1-ts* and *cnp1*Δ cells and further supporting their role as functional CENP-A protein, *P. murina* CENP-A protein also colocalize with *cen2-tetO-tdTomato* and remarkably exhibited single foci, albeit the *P. murina* CENP-A protein also colocalized with *cen2-tetO-tdTomato* and, remarkably, exhibited single foci, albeit the intensities of centromere foci were dimmer than that of the *S. pombe* counterpart (**Figure 2F**). Finally, to confirm the specific localization of *Pm* CENP-A to centromeres, we performed ChIP-qPCR (**Figure 2G**). Noteworthy, we observed that *Pm* CENP-A localizes to central core regions (i.e., *cc1&3, cc2:ura4^+^*) similarly to the fission yeast counterpart, albeit the enrichment was lower consistent with the results from live-cell imaging. Additionally, *Pm* CENP-A localization seems to be specific as no detectable enrichment was found at pericentromeric outer repeats coated with heterochromatin (i.e., *dg*) and euchromatin locations (i.e., *fbp1*). Together, these results confirm that *P. murina* CENP-A gene encodes a bona fide CENP-A protein.

**Fig. 2.**
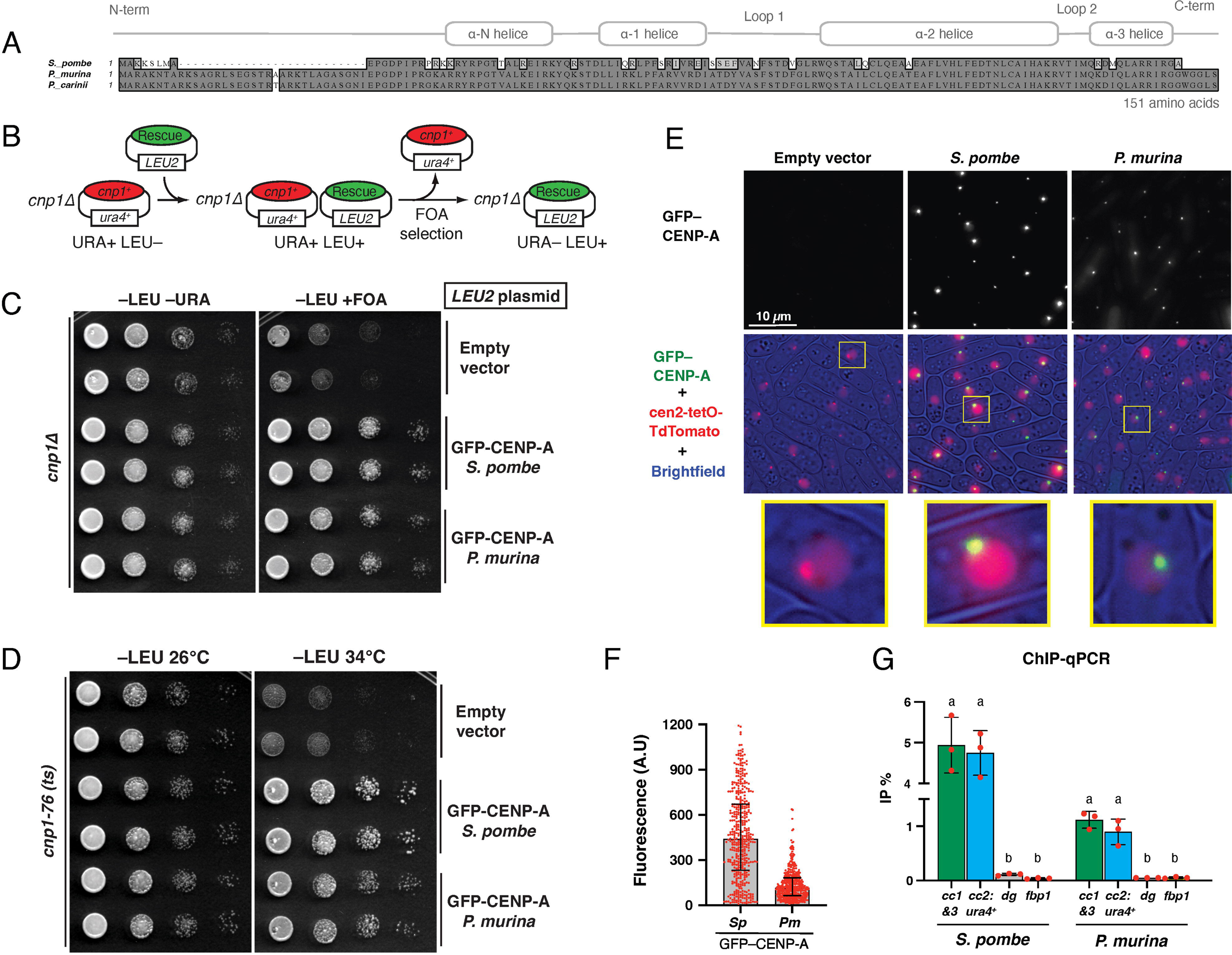
*Pneumocystis* CENP-A supports viability and centromere loading in a heterologous system. **(A)** Protein sequence alignment of full-length CENP-A^cnp-1^ orthologs in *Schizosaccharomyces pombe*, *Pneumocystis murina* and *P. carinii*. Each *Pneumocystis* species only infects a single mammalian host: *P. murina* (infecting mice) and *P. carinii* (rats). The ∼25 amino acid insert in *Pneumocystis* CENP-A N-terminal regions is unique to this genus. (**B**) Schematic of the plasmid-shuffling assay used in C. The inviability of *cnp1*Δ cells is masked by the presence of *cnp1^+^* on a *ura4^+^* marked plasmid. After introducing the *LEU2* marked rescue plasmid, cells that have lost the *ura4^+^* marked plasmid are selected on medium containing FOA, leaving only those with the rescue plasmid. NB: *S. cerevisiae LEU2* gene complements the *S. pombe leu1-32* mutant. (**C**) Rescue experiments using the plasmid-shuffling assay described in B to test the ability of multicopy plasmids expressing GFP-tagged CENP-A transgenes to rescue *cnp1*Δ cells. Growth was assayed at indicated media using 10-fold serial dilutions. (**D**) Rescue experiment of fission yeast *cnp1-76* thermosensitive cells using same plasmids as in C. Growth was assayed at indicated temperatures using 10-fold serial dilutions. The transgene encoding *S. pombe* CENP-A serves as a non-temperature-sensitive control. (**E**) Representative images of *cen2-tetO-TdTomato* strains expressing the indicated GFP-tagged CENP-A transgenes in multicopy plasmids. GFP and TdTomato fluorescence as well as brightfield images are merged. Magnified views of boxed regions are shown at the bottom. (**F**) Integrated fluorescence intensity of GFP foci was measured and plotted for the strains used in panel E. The median (bar) and interquartile range (error bars) are shown. n>409. AU, arbitrary units. (**G**) Anti-GFP ChIP-qPCR analysis of indicated loci for strains expressing GFP-tagged CENP-A transgenes as single copy integrations. %IP represents the percentage of input that was immunoprecipitated. Error bars denote the SD (n = 3). For each strain, comparison of %IP between loci was performed by one-way ANOVA and Holm-Sidak test for multiple comparisons (P<0.001). Mean values marked with letters (a or b) indicate results that are significantly different from each other. Abbreviations: *cc1&3*, *S. pombe* centromere central core 1 & 3; *cc2:ura4^+^*, *ura4*^+^ insertion at central core 2; *dg*, a class of heterochromatic outer repeats within pericentromeric regions (control); *fbp1*, euchromatic locus (control).

### *pn*CENP-A binds to single genomic foci in *Pneumocystis* replicating cells

In *S. pombe*, CENP-A^Cnp1^ is deposited at the centromeres during the G2 phase of the cell cycle (*31*). Although *Pneumocystis* cell replication is not fully understood, these organisms take 5 to 8 days to replicate *in vivo* (*36*). To determine *pn*CENP-A bound genome regions, we established a short-term co-culture system for *P. murina* and *P. carinii* (**Figure 3A**). We assessed organism growth using quantitative PCR targeting the single copy gene dihydrofolate reductase (*dhfr*); a typical growth curve shows a decline from day 0 to 7 followed by a gradual increase (**Figures 3B and 3C**). Population growth is further supported by the presence of mitotic or fusing cells 7 days post culture, though some cells do not appear to be dividing (**Figure 3D**).

**Fig. 3.**
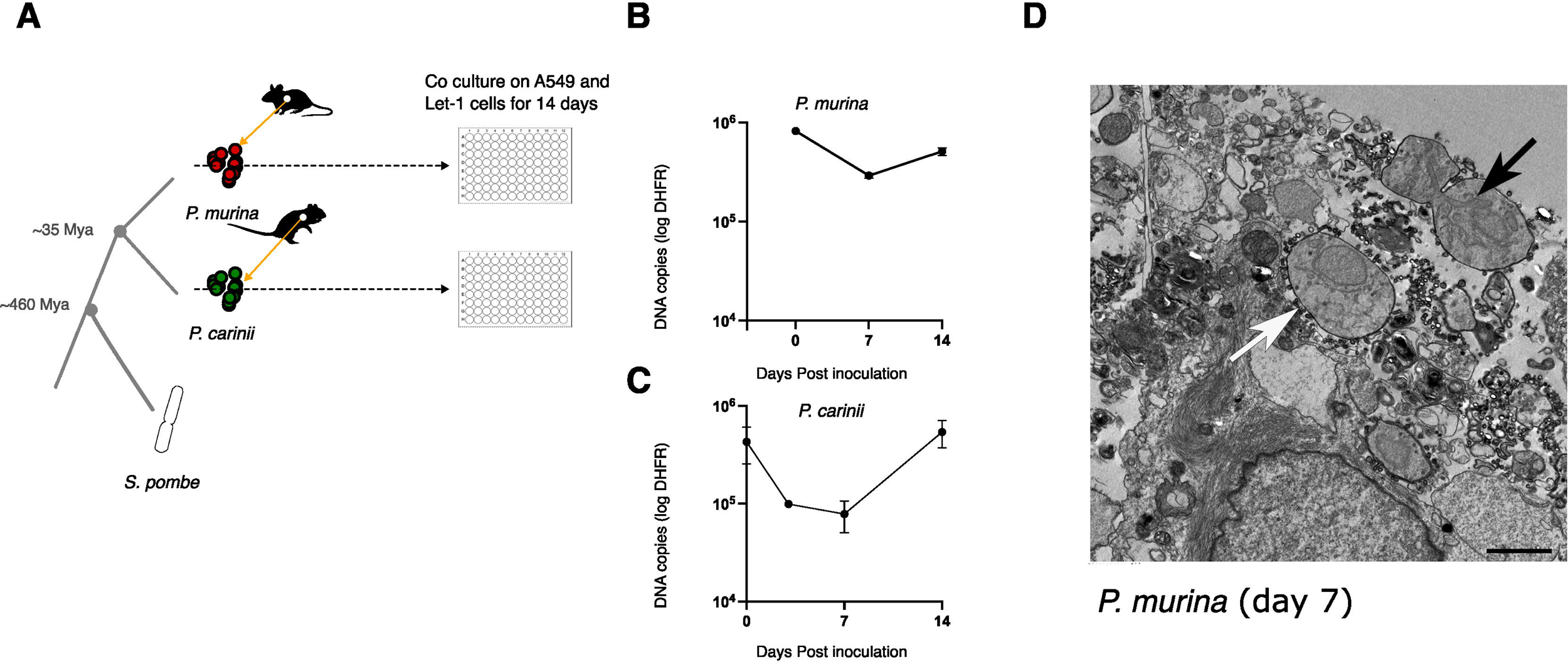
*Pneumocystis* CENP-A supports viability and centromere loading in a heterologous system. *Pneumocystis* CENP-A binds to single genomic foci in replicating cells. **(A)** Study workflow and sample preparation. *P. murina* and *P. carinii* were obtained from CD40 ligand knock-out female mice and immunosuppressed Sprague-Dawley male rats, respectively, and cultured on a co-culture of human lung adenocarcinoma cells (A549) and immortalized murine lung epithelial type 1 cells (Let-1) for 14 days. Species phylogeny and estimated speciation timing are presented. Animal icons were obtained from http://phylopic.org under creative commons licenses https://creativecommons.org/licenses/by/3.0/: mouse (Anthony Caravaggi; license CC BY-NC-SA 3.0); and rat (by Rebecca Groom; license CC BY-NC-SA 3.0). (**B**) *P. murina* population growth in duplicate wells were measured by quantitative PCR targeting a single copy dihydrofolate reductase (*dhfr*) gene over 14 days. Error bars represent the standard deviation (*n*=2). (**C**) *P. carinii* growth measurement by qPCR targeting *dhfr* gene. Error bars represent the standard deviation (*n*=2). (**D**) Electron micrograph showing a possibly dividing *P. murina* trophic form seven days post culture (black arrow). The field also include a non-dividing trophozoite (white arrow). 2 μm scale bar.

We mapped CENP-A bound genomic regions in all *P. carinii* and *P. murina* chromosomes using ChIP-from cultured organisms collected days 0 (normalized initial inoculum), 7 and 14.

In *P. murina*, each of the 17 chromosome-level scaffolds display a single region containing peaks of CENP-A at all three time points (**Figures 4A and 4B, Supplementary figure 4**). We confirmed CENP-A enrichment by ChIP-qPCR (**Supplementary figure 5A)**. CENP-A enrichment has a bimodal distribution, suggesting two distinct populations of CENP-A in chromosome arms. The two CENP-A peaks are separated by a 1-kb noncoding region that lacks a conserved DNA motif. Although the significance of two peaks of CENP-A in *Pneumocystis* is unclear, in *S. pombe* CENP-A cores are interspersed with H3K4me (*37*). CENP-A enrichment levels fluctuate in according to the growth curve, that is the highest level is observed at day 0 followed by a decline at day 7 and a recovery at day 14. In *S. pombe*, CENP-A^Cnp1^ enrichment at the centromeres correlates with a depletion of histone H3 (*38*). At day 7, the H3 occupancy (expressed as ratio between H3 and H4) is significantly reduced at centromeres relative to flanking regions (30 kb) as estimated by ChIP-Seq (Wilcoxon test; *P* < .0001; **Supplementary figure 6**), which suggest that CENP-A substitutes the canonical H3. However, H3 depletion at the centromeres does not persist after 14 days of culture (**Supplementary figure 7A**).

**Fig. 4.**
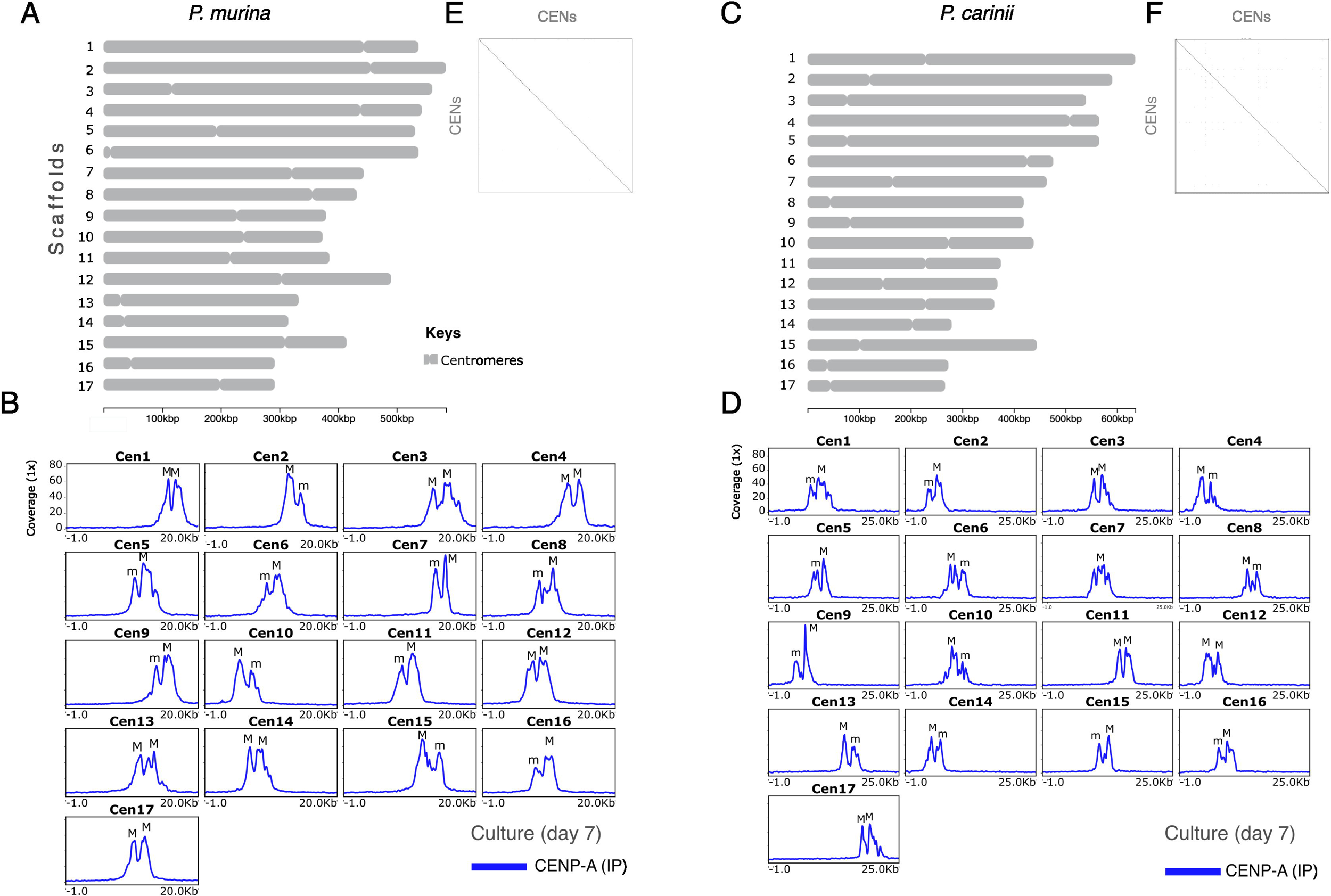
*Pneumocystis* displays seventeen centromeres. **(A)** Scaled ideogram of 17 chromosomal level scaffolds (grey) of *P. murina* genome showing CENP-A binding regions (constricted areas). (**B**) CENP-A binding regions delineate putative centromeres (CENs) in *P. murina* genome. Color coded peaks represent enrichment of immunoprecipitated DNA (IP DNA) relative to controls (Input DNA) in *P. murina* organisms (input subtracted). Only ChIP-Seq data from *Pneumocystis* cocultured with A549 and Let-1 cells for seven days are presented. Data for the full experiment covering days 0, 7 and 14 are presented in Supplementary material. The 20-kb windows of the enrichment peak are presented. Each scaffold displays two peaks labelled M (Major) and m (minor) according to the enrichment level. (**C**) Scaled ideogram of 17 chromosomal level scaffolds (grey) of *P. carinii* genome showing CENP-A binding regions (constricted areas). (**D**) CENP-A binding regions delineate putative centromeres (CENs) in *P. carinii* genome. Color coded peaks represent enrichment of immunoprecipitated DNA (IP DNA) relative to controls (Input DNA) in *P. carinii* organisms cocultured with A549 and Let-1 cells for seven days (input subtracted). Data for the full experiment covering days 0, 7 and 14 are presented in Supplementary figure 6 and Supplementary dataset. (**E**)A DNA dot plot of CENP-A binding regions in *P. murina* showing regional self-similarity. The main diagonal represents the sequence alignment with itself. Lines off the main diagonal which are repetitive patterns within the sequences are not observed. The plot shows that each CEN is unique within the genome. (**F**) DNA dot plot of CENP-A binding regions in *P. carinii* showing that each CEN is unique and repeat-free.

*P. carinii* also displays monocentric chromosomes enriched with two CENP-A peaks at the three time points by ChIP-Seq (**Figures 4C and 4D, Supplementary figure 8**) and ChIP-qPCR (**Supplementary figure 5B**). H3 depletion is observed at day 14 (**Supplementary figure 7B**). In contrast to *P. murina*, we found a significant reduction of CENP-A enrichment at day 14 despite a population growth (**Figure 3C**). Also not observed in *P. murina*, at day 14 low levels of CENP-A are present outside the primary CENP-A binding region in several chromosomes (**Supplementary figure 8**), which correspond to a non-specific displacement of CENP-A. In budding yeasts and chicken cells, low levels of CENP-A molecules have been shown outside the core domain (CENP-A cloud), which contribute to centromere plasticity (*39*).

In both species CENP-A bound regions span 4.8 to 8.0 kb, with an average length of 6.7 kb, and have an average of 3.9% lower GC content than the rest of the genome (**Table 1**). Using dot plot analysis of a concatemer of different centromeres, each CENP-A binding region appears unique within the genome (**Figures 4E and 4F**) and lacks a shared conserved DNA motif (see methods). Because most centromeres in fungi are associated with repetitive DNA (e.g., DNA transposons and retrotransposons), we searched for known signatures of repeats and found no significant overlap of repeats with centromeres (**Supplementary figures 6 and 9**).

**Table 1.**
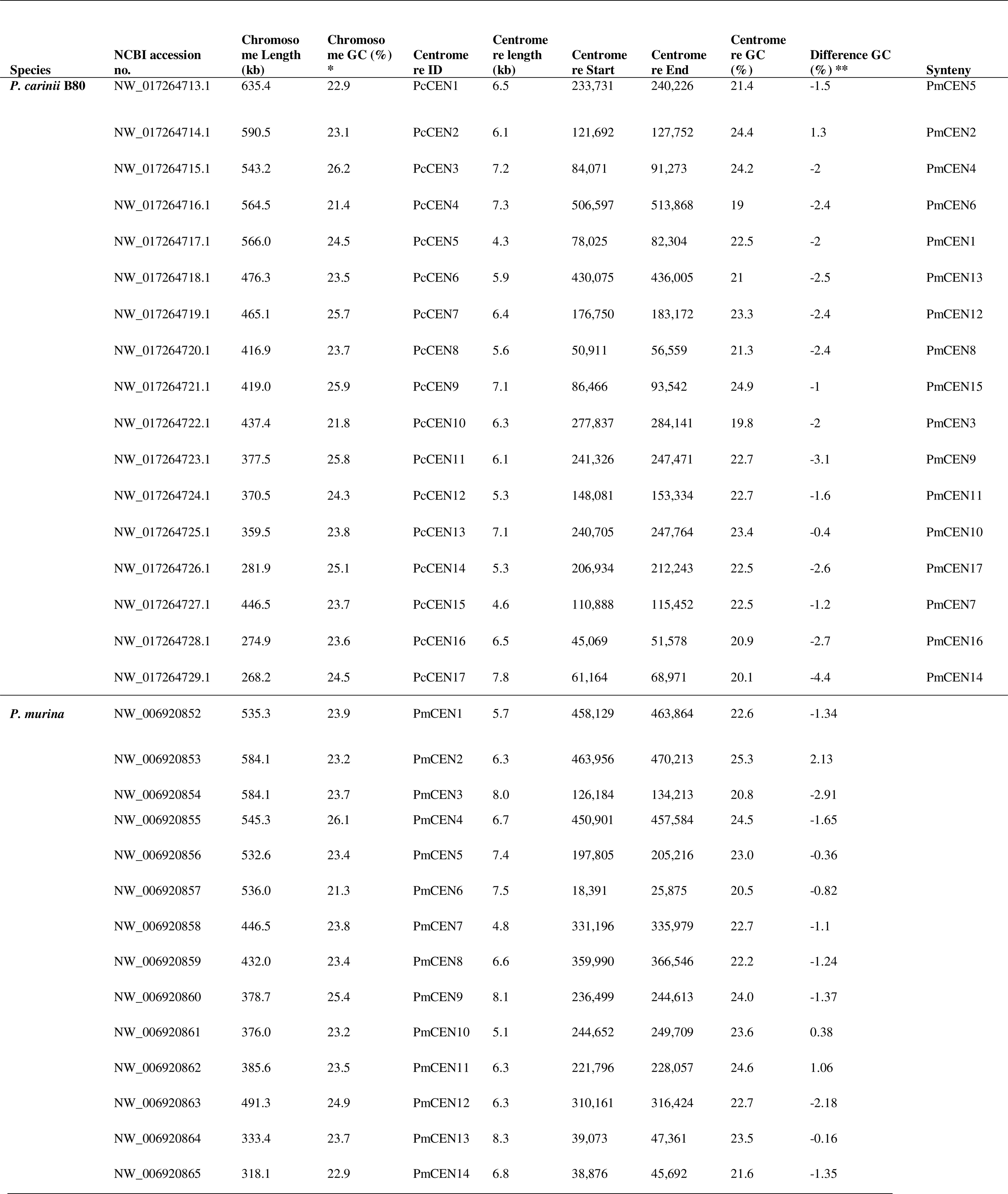
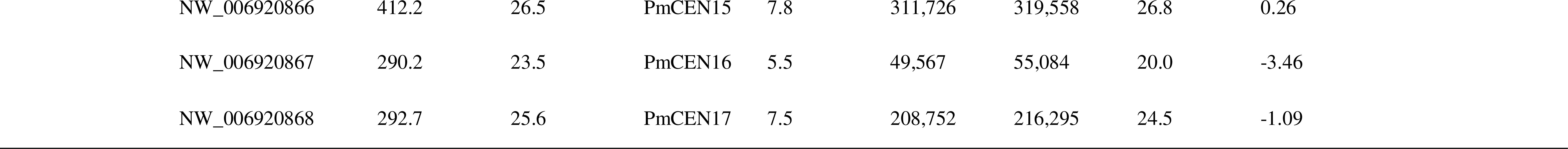
Coordinates, length, and GC content (in %) of *Pneumocystis* centromeres identified by direct ChIP.

| Species | NCBI accession no. | Chromosome Length (kb) | Chromosome GC (%) * | Centromere ID | Centromere length (kb) | Centromere Start | Centromere End | Centromere GC (%) | Difference GC (%) ** | Synteny |
| --- | --- | --- | --- | --- | --- | --- | --- | --- | --- | --- |
| *P. carinii* B80 | NW_017264713.1 | 635.4 | 22.9 | PcCEN1 | 6.5 | 233,731 | 240,226 | 21.4 | -1.5 | PmCEN5 |
|  | NW_017264714.1 | 590.5 | 23.1 | PcCEN2 | 6.1 | 121,692 | 127,752 | 24.4 | 1.3 | PmCEN2 |
|  | NW_017264715.1 | 543.2 | 26.2 | PcCEN3 | 7.2 | 84,071 | 91,273 | 24.2 | -2 | PmCEN4 |
|  | NW_017264716.1 | 564.5 | 21.4 | PcCEN4 | 7.3 | 506,597 | 513,868 | 19 | -2.4 | PmCEN6 |
|  | NW_017264717.1 | 566.0 | 24.5 | PcCEN5 | 4.3 | 78,025 | 82,304 | 22.5 | -2 | PmCEN1 |
|  | NW_017264718.1 | 476.3 | 23.5 | PcCEN6 | 5.9 | 430,075 | 436,005 | 21 | -2.5 | PmCEN13 |
|  | NW_017264719.1 | 465.1 | 25.7 | PcCEN7 | 6.4 | 176,750 | 183,172 | 23.3 | -2.4 | PmCEN12 |
|  | NW_017264720.1 | 416.9 | 23.7 | PcCEN8 | 5.6 | 50,911 | 56,559 | 21.3 | -2.4 | PmCEN8 |
|  | NW_017264721.1 | 419.0 | 25.9 | PcCEN9 | 7.1 | 86,466 | 93,542 | 24.9 | -1 | PmCEN15 |
|  | NW_017264722.1 | 437.4 | 21.8 | PcCEN10 | 6.3 | 277,837 | 284,141 | 19.8 | -2 | PmCEN3 |
|  | NW_017264723.1 | 377.5 | 25.8 | PcCEN11 | 6.1 | 241,326 | 247,471 | 22.7 | -3.1 | PmCEN9 |
|  | NW_017264724.1 | 370.5 | 24.3 | PcCEN12 | 5.3 | 148,081 | 153,334 | 22.7 | -1.6 | PmCEN11 |
|  | NW_017264725.1 | 359.5 | 23.8 | PcCEN13 | 7.1 | 240,705 | 247,764 | 23.4 | -0.4 | PmCEN10 |
|  | NW_017264726.1 | 281.9 | 25.1 | PcCEN14 | 5.3 | 206,934 | 212,243 | 22.5 | -2.6 | PmCEN17 |
|  | NW_017264727.1 | 446.5 | 23.7 | PcCEN15 | 4.6 | 110,888 | 115,452 | 22.5 | -1.2 | PmCEN7 |
|  | NW_017264728.1 | 274.9 | 23.6 | PcCEN16 | 6.5 | 45,069 | 51,578 | 20.9 | -2.7 | PmCEN16 |
|  | NW_017264729.1 | 268.2 | 24.5 | PcCEN17 | 7.8 | 61,164 | 68,971 | 20.1 | -4.4 | PmCEN14 |
| *P. murina* | NW_006920852 | 535.3 | 23.9 | PmCEN1 | 5.7 | 458,129 | 463,864 | 22.6 | -1.34 |  |
|  | NW_006920853 | 584.1 | 23.2 | PmCEN2 | 6.3 | 463,956 | 470,213 | 25.3 | 2.13 |  |
|  | NW_006920854 | 584.1 | 23.7 | PmCEN3 | 8.0 | 126,184 | 134,213 | 20.8 | -2.91 |  |
|  | NW_006920855 | 545.3 | 26.1 | PmCEN4 | 6.7 | 450,901 | 457,584 | 24.5 | -1.65 |  |
|  | NW_006920856 | 532.6 | 23.4 | PmCEN5 | 7.4 | 197,805 | 205,216 | 23.0 | -0.36 |  |
|  | NW_006920857 | 536.0 | 21.3 | PmCEN6 | 7.5 | 18,391 | 25,875 | 20.5 | -0.82 |  |
|  | NW_006920858 | 446.5 | 23.8 | PmCEN7 | 4.8 | 331,196 | 335,979 | 22.7 | -1.1 |  |
|  | NW_006920859 | 432.0 | 23.4 | PmCEN8 | 6.6 | 359,990 | 366,546 | 22.2 | -1.24 |  |
|  | NW_006920860 | 378.7 | 25.4 | PmCEN9 | 8.1 | 236,499 | 244,613 | 24.0 | -1.37 |  |
|  | NW_006920861 | 376.0 | 23.2 | PmCEN10 | 5.1 | 244,652 | 249,709 | 23.6 | 0.38 |  |
|  | NW_006920862 | 385.6 | 23.5 | PmCEN11 | 6.3 | 221,796 | 228,057 | 24.6 | 1.06 |  |
|  | NW_006920863 | 491.3 | 24.9 | PmCEN12 | 6.3 | 310,161 | 316,424 | 22.7 | -2.18 |  |
|  | NW_006920864 | 333.4 | 23.7 | PmCEN13 | 8.3 | 39,073 | 47,361 | 23.5 | -0.16 |  |
|  | NW_006920865 | 318.1 | 22.9 | PmCEN14 | 6.8 | 38,876 | 45,692 | 21.6 | -1.35 |  |

|  |  |  |  |  |  |  |  |  |
| --- | --- | --- | --- | --- | --- | --- | --- | --- |
| NW_006920866 | 412.2 | 26.5 | PmCEN15 | 7.8 | 311,726 | 319,558 | 26.8 | 0.26 |
| NW_006920867 | 290.2 | 23.5 | PmCEN16 | 5.5 | 49,567 | 55,084 | 20.0 | -3.46 |
| NW_006920868 | 292.7 | 25.6 | PmCEN17 | 7.5 | 208,752 | 216,295 | 24.5 | -1.09 |

### *Pneumocystis* centromeres contain active genes

By cross referencing annotated gene locations with those of *pn*CENP-A bound regions, we found that *P. murina* and *P. carinii* centromeres encode 74 and 58 genes respectively; all of them are conserved in the genomes of both species except a prefoldin gene that is lost in *P. carinii* (**Supplementary Table 2**). Of the 74 *P. murina* centromeric genes, 73 were expressed based on RNA-seq and four were further detected by protein mass spectrometry mapping (LC-MS); of the 58 *P. carinii* genes, 56 are expressed and 53 were detected by LC-MS. Analysis of the predicted function of these genes revealed housekeeping functions without major differences compared to randomly sampled genes.

### Centromeres are flanked by heterochromatin

In *S. pombe*, H3K9me2/3 marking of chromatin is associated with the repression of centromeres, subtelomeres, ribosomal rDNA and the mating locus (*37*). There are two types of chromatins: euchromatin, the lightly packed form of the chromatin enriched with H3K4me2 (di-methylation of the 4^th^ lysine residue of the histone H3), which is associated with active transcription, and heterochromatin enriched with H3K9me2 and H3K9me3 (di and tri-methylation of the 9^th^ lysine residue of the histone H3 protein). To test if this feature is shared in *Pneumocystis*, we performed ChIP-Seq with antibodies targeting histones H3 modifications (H3K9me2/3 and H3K4me2). In *Pneumocystis*, we detected broadly distributed peaks of H3K9me2 and H3K9me3 bordering the putative centromeres delineated by CENP-A narrow peaks (**Figures 5A and 5B**, **Supplementary figures 6 and 9**). This configuration of chromatin markers is specific to centromeric regions. H3K4me2 is correlated with H3K9me2 in the centromeres (Pearson rho = .72) but not at the whole genome level (r = .05). However, research has shown that the histone modifications H3K4me2 and H3K9me2 decorate euchromatin and heterochromatin regions, respectively (*40*). Therefore, the correlation observed between H3K4me2 and H3K9me2 may be due to differences in growth kinetics for the *Pneumocystis* population of cells used in the ChIP analysis. Furthermore, similar to *S. pombe* (*37*), a small amount of H3K4me2 is present at the centromeres in *Pneumocystis* (**Figures 5A and 5B**, **Supplementary figures 6 and 9**). These results suggest that *Pneumocystis* centromeres are flanked by high levels of heterochromatin.

**Fig. 5.**
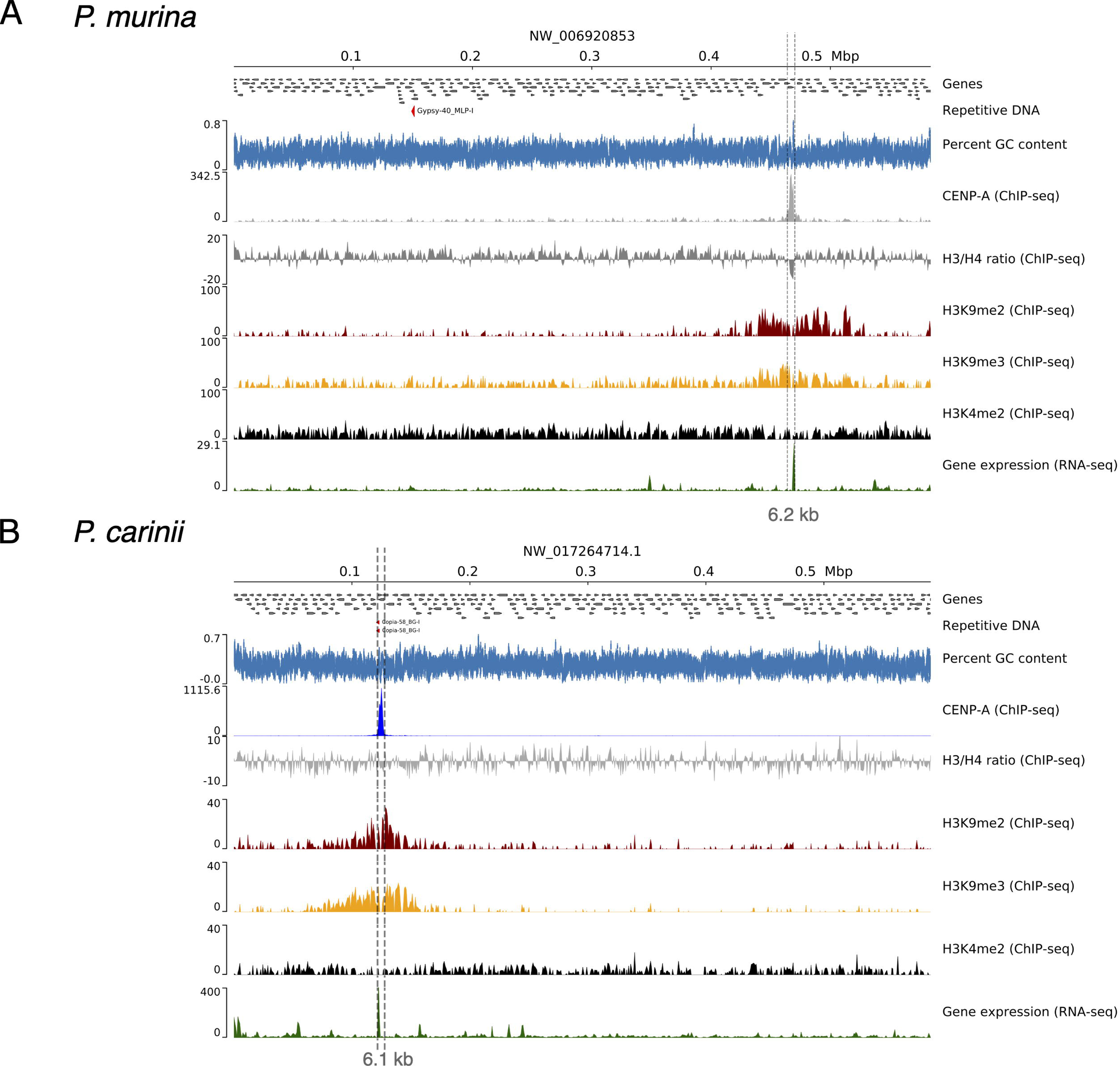
Centromeres are flanked by heterochromatin and contain active genes. (**A**) Genomic view of chromosome 2 of *P. murina* genome subsequently showing annotated genes (directed grey boxes), DNA repeats (not present), percent GC content (blue), ChIP-Seq read coverage distribution (BPM normalized over bins of 50 bp; input subtracted) of CENP-A, histones H3 and H4 ratio, heterochromatin-associated modifications (H3K9me2 and H3K9me3), euchromatin (H3K4me2) and gene expression (RNA-seq) in relation with centromeres. (**B**) Genomic view of chromosome 2 of *P. carinii* genome with the same features presented as *P. murina*. A duplicated copy of *copia*-retrotransposon is presented (red arrows), which is present in syntenic regions in other *Pneumocystis* genomes. The two presented chromosomes are syntenic.

DNA methylation is frequently associated with centromeres in fungi, where they silence repeats (*41, 42*). Similar to *S. pombe*, DNA methylation has been predicted to be absent in *Pneumocystis* based on the absence of DNA methyltransferases (DMTs) (*29, 43*). To determine whether DNA methylation plays a role in *Pneumocystis* centromeres, we performed bisulfite sequencing (5 methylcytosine) in *P. carinii*. The overall level of 5mC DNA methylation as measured by the average weighted methylation percentage is 0.6% for *P. carinii* at the CG dinucleotides (**Supplementary figure 10**). These levels are in range with reported levels for other fungi e.g., *Verticillium* (0.4%) (*44*). The presence of 5mC methylated DNA bases despite the absence of recognizable DNA methylases in *Pneumocystis* requires further investigation. To assess the potential role of DNA methylation in centromere function in *Pneumocystis*, we analyzed the DNA methylation patterns over different genomic features (genes, intergenic spacers, and centromeres). However, there is no significant difference among centromeric or pericentromeric regions (defined as 30 kb flanking the centromeres) (Mann-Whitney U-test *P*-value > 0.3), and the randomly selected genomic regions (genomic background). These results suggest that 5-mC DNA methylation is not required for centromere function in *Pneumocystis*. The absence of repeats in the centromeres and the presence of actively transcribed genes suggests that DNA methylation does not induce an epigenetic silencing in *Pneumocystis* centromeres.

### Centromere conservation during speciation

*Pneumocystis murina and P. carinii* species have diverged ∼ 35 Mya (*12*). Yet, orthologous regions act as centromeres in both *species,* which suggest a vertical transmission (**Figure 6A**). Syntenic regions can be found across five additional *Pneumocystis* species which speciation spans at least 100 million years (**Supplementary figure 11**).

**Fig. 6.**
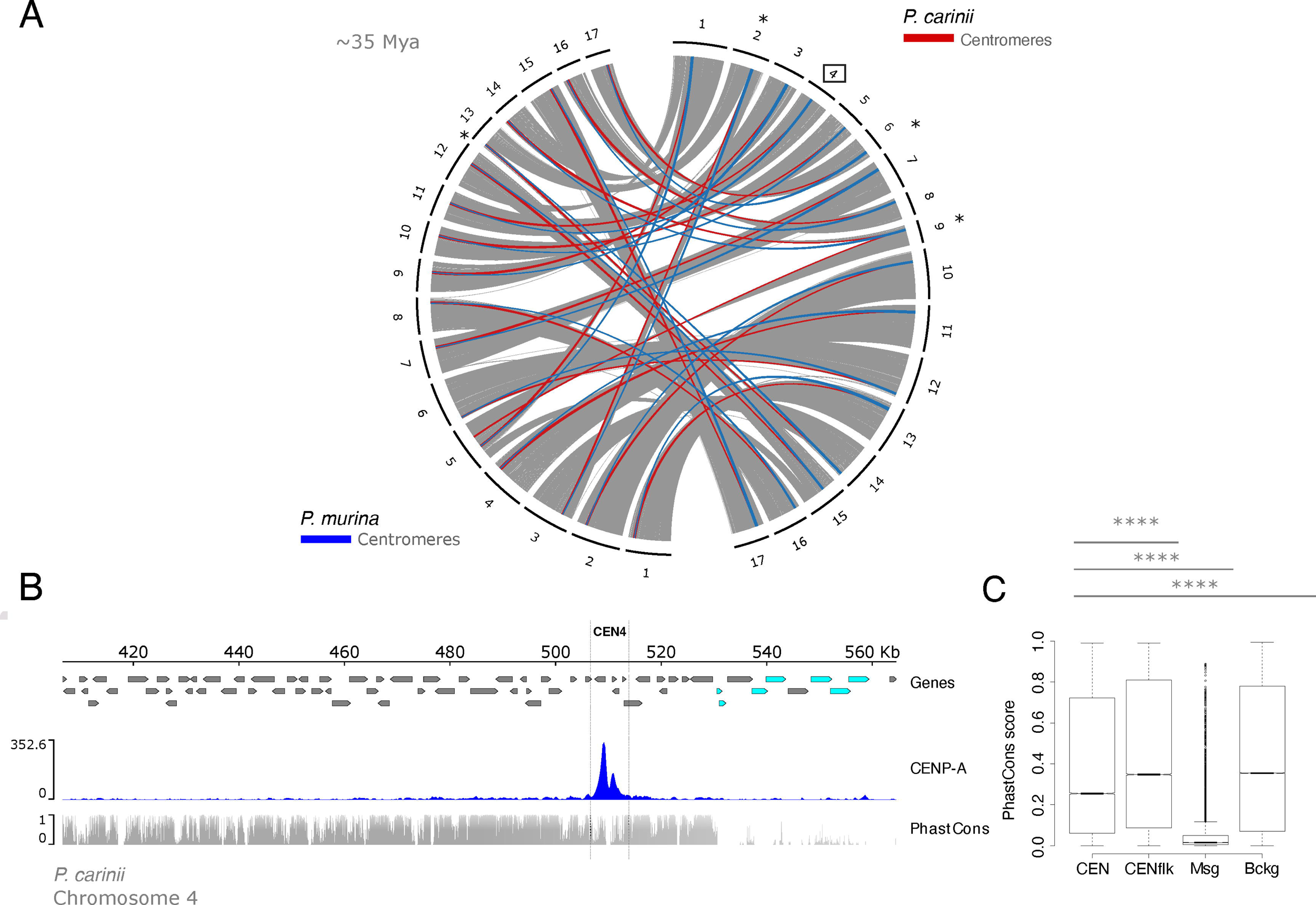
Centromere locations, not sequences, are evolutionarily constrained by positive selection. **(A)** Genome organization and synteny in *Pneumocystis*. Circos plots depicting pairwise *P. carinii* and *P. murina* genome synteny. Rodent infecting *Pneumocystis* (*P. carinii* and *P. murina*) have 17 chromosomes. Colored connectors indicate regions of synteny between species. Centromeres that overlap with recent chromosomal breakpoints are indicated (*). The square highlights *P. carinii* centromere 4 displayed in the panel b. (**B**) Genome view of centromere 4 in the *P. carinii* genome. Genes are represented by directed boxes (grey for protein coding genes and cyan for polymorphic major surface glycoprotein genes), *pn*CENP-A binding region (centromere) and sequence conservation scores which were calculated from whole genome alignments of *P. carinii*. *P. murina* and *P. wakefieldiae* (PhasCons). The phastCons scores represent probabilities of negative selection and range between 0 (no conservation) and 1 (total conservation). (**C**) Boxplot of conservation scores per genomic context summarized for the centromere 4 in *P. carinii* genome, 30-kb regions flanking the centromeres (Cenflk), major surface glycoproteins encoding regions (Msg) and random genomic background (Bckg). Msgs are fast evolving proteins potentially involved in antigenic variation. Background data were obtained from randomly selected intervals (*n* = 1 x 10^6^) from genomic regions excluding above mentioned regions (CEN, Cenflk and Msg). Statistical differences for the indicated comparisons were obtained using one-sided nonparametric Mann-Whitney test; *P*-values < .0001, ****; *P* < .001, ***; *P* < .01, **; *P* < .05, *. Data for all 17 centromeres are presented in Supplementary material.

To investigate footprints of selection acting on centromeres, we computed conservation scores (PhastCons) from whole genome alignments. Conservation scores range from 0 to 1 and represent probabilities of negative selection. *Pneumocystis* genomes encode a large multicopy major surface glycoprotein (MSG) gene family. MSGs are highly polymorphic and evolve rapidly due to their role in antigenic variation during mammalian host infection (reviewed in (*45*)). Using MSGs as control for evolutionary speed, we found that centromeres and flanking regions are substantially more conserved than MSGs and similar to genomic background (**Figures 6B and 6C**). The levels of centromeric conservation are variable across the chromosomes, with the flanking regions being more conserved than the cores (defined as a 1-2 kb region in the middle of each centromere) (**Supplementary figure 12)**. Given that macrosynteny at the centromeric regions is conserved across multiple *Pneumocystis* species, this suggests that centromere locations are maintained by positive selection. This would be consistent with the hypothesis that centromere positioning tends to be conserved in obligate sexual fungi due to their role in meiosis (*46*).

Centromeres contribute to karyotypic diversity in some fungi (*2*). However, we found little support for this hypothesis here because most centromeres do not overlap with chromosomal breaks (only 3 of 17 in the *P. carinii* versus *P. murina* pairwise comparison). This is reminiscent of other fungi such as *Verticillium* species where centromeres do not account for most of the karyotype variation (*47*).

## Discussion

In the current study we have identified centromeres of two *Pneumocystis* species by showing that the centromeric histone CENP-A binds to gene rich genomic regions that are flanked with heterochromatin (**Figure 7**).

**Fig. 7.**
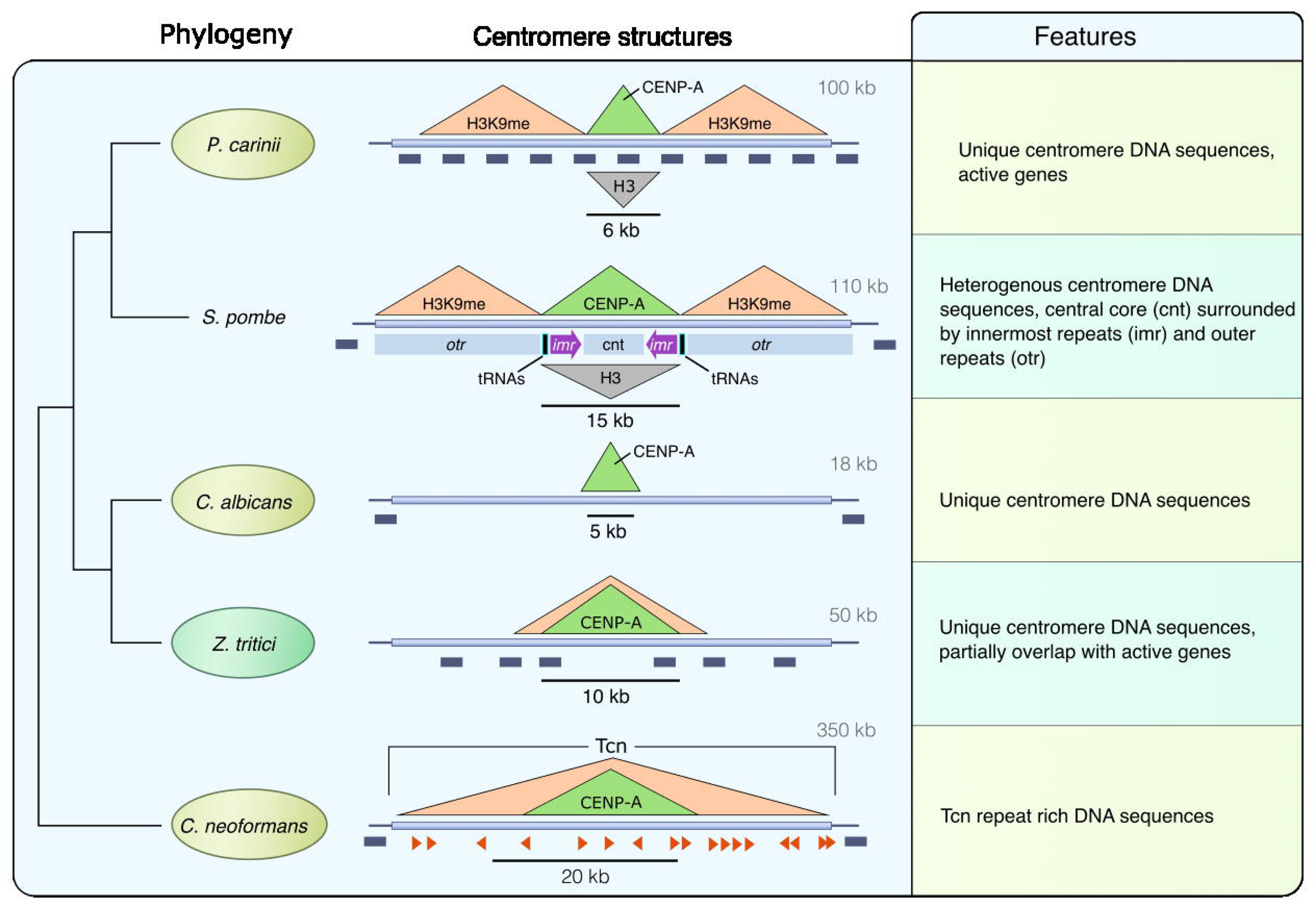
Model of regional centromere structures in *Pneumocystis* and other representative fungi. On the left is presented the phylogeny of selected fungi inferred from maximum likelihood phylogenetic analysis of shared core protein orthologs. The overall centromere structures for *Pneumocystis carinii* (details supporting our model are provided in the text), *Schizosaccharomyces pombe* (*48*), *Candida albicans* (*8*), *Zymoseptoria tritici* (*46*) and *Cryptococcus neoformans* (*49*) are used as representative to showcase the diversity of regional centromeres in fungi. Animal pathogens are highlighted in pale olive and the plant fungal pathogen *Zymoseptoria tritici* is in pale green. For sake of brevity, only *Pneumocystis carinii* is presented. In the middle are presented DNA structures of centromeres. *P. carinii* has 17 centromeres (one for each of its 17 chromosomes) that share the same overall architecture. Centromeres are delineated by a localized enrichment of the centromeric histone CENP-A, which overlaps with a reduction of the canonical histone H3 (inverted grey triangle). Centromeres are flanked by heterochromatin H3K9me (stands here stands for both H3K9me2 and H3K9me3). All 17 *Pneumocystis carinii* centromeres span active genes (dark grey boxes). Each centromere sequence is different and lacks shared DNA sequence motif. *S. pombe* has three centromeres that share the same overall structure in which a central core (*cnt*) domain is surrounded by innermost repeats (*imr*) and outer repeats (*otr*). The *imr* repeats incorporate clusters of transfer RNAs (tRNAs) that play a role in restricting CENP-A spread. *S. pombe* centromeres are flanked by heterochromatin (H3K9me). Genes are found 0.75-1.5 kb beyond the limits of the centromeres. *C. albicans* has 8 unique and different centromeres that are gene free and lack shared sequence motifs. *Z. tritici* has 21 centromeres ranging from 6 to 14 kb in size, that partially overlap with genes. *C. neoformans* has 14 centromeres that are gene free and enriched with Tcn transposons.

This configuration suggests the presence of small epigenetically regulated centromeres (regional). Within each genome, each chromosome has a unique non repetitive centromere. Syntenic regions for each centromere are present in all seven currently sequenced *Pneumocystis* species. Cell replication and chromosomal segregation pathways are unexplored in these species. Central to these pathways are the centromeres.

As centromeres cannot be predicted bioinformatically because no suitable reference for synteny analyses existed before this study, the locations and characteristics of *Pneumocystis* centromeres were unknown. Moreover, there are several roadblocks for characterizing centromeres in *Pneumocystis*, with the most significant one being the lack of continuous culture and transfection tools. Here we utilized two *Pneumocystis* species in a short-term culture system to determine where *pn*CENP-A, the epigenetic marker for centromeres, binds in the genome.

As a first step, we demonstrated that *Pn*CENP-A functions similarly to CENP-A*^Cnp1^* in *S. pombe*, though less efficiently, validating its use in characterizing *Pneumocystis* centromeres. We then developed a *Pn*CENP-A based ChIP-Seq assay that showed a reproducible efficiency in defining centromeric regions of *Pneumocystis* species.

Unlike the phylogenetically related *S. pombe*, in *Pneumocystis* these regions overlap with active genes, but like *S. pombe*, these centromeres are flanked by heterochromatin. Our work also provides the first experimental evidence that DNA methylation occurs in these species, although it may not be involved in centromere function.

Despite their close phylogenetic relationship, *P. murina* and *P. carinii* have slightly different growth kinetics in our short-term culture system, which could explain the variations in the *Pn*CENP-A and H3 DNA binding profiles. The nonspecific binding of *Pn*CENP-A observed in *P. carinii* at day 14 is likely the result of a random dissociation of CENP-A molecules from the centromeres.

In *P. murina*, 5 out of the 17 chromosomes showed secondary CENP-A enrichment at non centromeric sites (Figure S4). These secondary CENP-A peaks are likely ChIP-Seq artefacts because they were only detected at day 7 post culture, and mostly overlap with highly expressed loci (Figure S8), which are susceptible to misleading signals in ChIP-Seq experiments (*50*).

The presence of genes within centromeres is rare and has only been described in rice (*51*) and the plant fungal pathogen *Zymoseptoria tritici* (*46*). Centromeres bearing genes are interpreted as young centromeres (neocentromeres), in which the genes are progressively inactivated. This hypothesis lacks support here because there is no sign of pseudogenization in these genes: nearly all are expressed and translated, they are conserved in all species, and many are involved in housekeeping cellular pathways that are presumably critical for organism survival.

Findings in *Pneumocystis* cannot be generalized to other Taphrinomycotina species. The determination of ancestral traits between *Pneumocystis* and fission yeast centromeres will require characterizing the centromeres in additional Taphrinomycotina species. The loss of RNAi is often associated with shortening of centromeres (*49*); consistent with this, *Pneumocystis* centromeres are much smaller than *Schizosaccharomyces* centromeres.

Our study will benefit from a microscopy validation of CENP-A loading kinetics when a reliable long term culture system becomes available. Live imaging would help to determine at which cell cycle stage CENP-A is loaded. A direct genetic confirmation of *Pneumocystis* centromeres will also be required when a long-term culture system and genetic manipulation tools will become available.

In summary, we have identified short regional centromeres in genetically intractable micro-organisms. Our results provide insights into the formation of centromeres in host-adapted fungal pathogens. Our ultimate goal is to use *Pneumocystis* centromeres to stabilize plasmids for future genetic manipulation. This is the first step along this path, which should lead to better understanding of the biology of *Pneumocystis* and facilitate the discovery of novel interventions to effectively control and prevent the disease caused by this pathogen.

## Materials and Methods

### Ethics and organism source

Studies involving mouse and rat samples were approved by the NIH Institutional Review Board (IRB) (IRB protocols 99-I-0084 and CCM 19-05, respectively), and rat *Pneumocystis* studies by the Animal Care and Use Committees of the Cincinnati VA Medical Center (protocol 20-11-08-01).

*P. carinii* organisms were collected from heavily infected lungs of corticosteroid-treated Sprague-Dawley male rats; *P. murina* from heavily infected lungs of CD40 ligand knock out mouse. The list of samples is presented in the **Supplementary table 3**.

### *Pneumocystis* culture and growth quantification

*P. murina* and *P. carinii* were partially purified by Ficoll-Hypaque density gradient centrifugation (*52*) and frozen at -80°C in cell recovery media (Gibco). For short term *Pneumocystis* cultures, A549 (ATCC) and LET1 (a gift from Dr. Paul Thomas, St Jude Children’s Research Hospital) cells were cultured in culture medium F12 media with 2.5% or 5% heat inactivated fetal bovine serum (GIBCO) and Penicillin-Streptomycin (GIBCO), plated to approximately 60 – 80% confluency and incubated for 24h at 37°C. The next day, frozen *Pneumocystis* vials were thawed, washed in 50ml 1x PBS, and centrifuged at 2,000g for 20 minutes. *Pneumocystis* cell pellets were resuspended in culture medium and added to the plated cells. Media were partially changed every 3 or 4 days. At set time points (days 7 and 14), wells were scraped for collection. Cell/organism suspensions were centrifuged at 10,000g for 3 min, the supernatant was removed, and the pellets were frozen at -80°C. Genomic DNA was extracted using QiAmp DNA Extraction Kit (Qiagen). Quantitative PCR targeting the single copy *dhfr* gene was performed as described previously (*53*).

### *P. murina* CENP-A^Cnp1^ complementation and ChIP-qPCR in *S. pombe*

Standard procedures were used for fission yeast growth, genetics, and manipulations (*54*). The sequence encoding *P. murina* CENP-A gene was codon optimized for *S. pombe* codon bias, synthesized commercially (GenScript, NJ) and subcloned into pS2 vector (*55*) generating pPmCENP-A. *Nde*I-*Sph*I fragments from both pS2 and pPmCENP-A were subcloned into the same sites of pREP81 vector to obtain *LEU2*-based multicopy plasmids that were used in experiments presented in Fig. 2B-F. Alternatively, plasmids were linearized with *Nsi*I and introduced by transformation to the *lys1^+^* locus of *cen2-tetO-tdTomato* strain (*56*). These strains bearing single-copy integrations were grown overnight at 30°C in rich medium YEA and used for ChIP with 2 µl of anti-GFP (ab290, Abcam). PCR oligonucleotides detecting central core (*cc1&3; ura4*), heterochromatic *dg* and euchromatin control *fbp1* were previously reported (*57*).

### Orthologs identification

To find orthologs for kinetochore proteins, we constructed a database of 21 proteomes from fungi 15 Taphrinomycotina including *Pneumocystis* (*n* = 7) and 6 other representative fungi (**Supplementary Table 1**). We queried the database with *S. pombe* proteins as seed using BLASTp (*58*) with a minimum an e-value of 1.0 x 10^-5^ as threshold.

If many homologs were detected in *Pneumocystis* species, we separated orthologs from out-paralogs by searching in orthologous groups inferred from the clustering of the whole proteome dataset using OrthoFinder v2.5.2 (*59*). A sequence was considered as ortholog if it was assigned to the same orthologous group as *S. pombe* protein query. We verified that the assignment was consistent with EggNOG database v6.0 (*60*), which contains all *S. pombe* proteins and a limited number of *Pneumocystis* proteins. If a *Pneumocystis* sequence was detected by BLASTp but not assigned as an ortholog, we first identified conserved domains in all sequences using InterProScan version 5.64-96.0 (*61*). Then we constructed multiple sequence alignment containing all sequences including *S. pombe* query using MAFFT version v7.467 (*62*) either with the option E-INS-I or L-INS-i depending on the protein domain architecture. Alignments were manually curated using MacVector version 18.6.1 (https://macvector.com/) to remove spurious aligned regions and used to infer an approximately maximum likelihood gene tree using FastTree v2.1.10 (*63*). When necessary, a maximum likelihood phylogeny was performed using IQ-TREE version 1.5.5 (*64*). For histones, we used protein sequences identified by (*65*) to guide the classification. In case of inconclusive phylogenetic classification, we BLASTed *Pneumocystis* protein sequences against NCBI nr using a minimum e-value of 1.0 x 10^-5^ as threshold. Retrieved proteins were added to the existing alignment, and the phylogenetic inference was repeated.

If no homologs were found in *Pneumocystis* using *S. pombe* query, we used a different seed from another *Taphrinomycotina* if available (e.g., *Saitoella*). Alternatively, proteins from, *Saccharomyces cerevisiae*, *Neurospora crassa*, and *Cryptococcus neoformans* were used as seeds. If no homologs were detected, we used PSI-BLASTp (*58*). In parallel, if *S. pombe* query contains a conserved protein domain, we screened our fungal database with the corresponding Pfam domain using hmmsearch (HMMER version 3.3.2; http://hmmer.org)(66) with a cut off e-value of 0.1.

Because many genes could not be found in *Pneumocystis* proteomes, we scanned the genomes using TBLASTn (BLAST+ 2.13.0)(*67*) with an e-value of 0.1 as threshold. If putative genomic regions were found, we performed using a protein-to-genome alignment with Exonerate (*68*). Protein to genome alignments were translated in protein sequences, aligned, and used phylogenetic inference as described above.

To identify DNA methylases within the Taphrinomycotina, we identified annotated proteins with a DNA methylase domain (Pfam domain PF000145(*69*)) using hmmsearch (HMMER version 3.3.2; http://hmmer.org)(66) with a cut off e-value of 0.01. Although no hits were found in *Pneumocystis* proteomes, DMTs were detected in other fungi including *Neolecta irregularis* and *Saitoella complicata*, which is consistent with a previous study (*28*). *Pneumocystis* proteomes were queried using sequences from *N. irregularis* and *S. complicata* using BLASTp (*58*) with an e-value of 1 x 10^-5^ as cut off. This strategy identified 52 *Pneumocystis* proteins with local similarities, which were combined with all DMTs identified, aligned using Muscle v3.8.31 (*70*) and used for phylogenetic inference using FastTree. Then, we BLASTed *Pneumocystis* proteins against NCBI nr database (last accessed Thu Oct 12 12:07:56 EDT 2023), retrieved the top hit sequences and added them to the multiple alignment. All 52 *Pneumocystis* sequences belong related but distinct protein families (e. g. DNA helicases and SNF2 chromatin remodeling ATPases).

### Electron microscopy

*P. murina* and A549-LET1 cells described above were grown on Thermanox™ coverslips (Ted Pella, Redding, CA, USA), harvested at day 7, fixed with 2% paraformaldehyde/2.5% glutaraldehyde in 0.1 M Sorenson’s phosphate buffer and then post-fixed with 1.0% osmium tetroxide/0.8% potassium ferricyanide in 0.1 M sodium cacodylate buffer for 1 hour, washed with buffer then stained with 1% tannic acid in dH_2_O for 1 hr. After additional buffer washes, the samples were further osmicated with 2% osmium tetroxide in 0.1M sodium cacodylate, then washed with dH_2_O. Specimens were then stained overnight at 4C with 1% aqueous uranyl acetate. The cells were then washed with dH_2_O and dehydrated with a graded ethanol series, prior to embedding in Spurr’s resin. Thin sections were cut with a Leica UC7 ultramicrotome (Buffalo Grove, IL) prior to viewing at 120 kV on a FEI BT Tecnai transmission electron microscope (ThermoFisher/FEI, Hillsboro, OR). Digital images were acquired with a Gatan Rio camera (Gatan, Pleasanton, CA).

### Antibodies

To obtain antibody against *pn*CENP-A, we created an alignment containing CENP-A orthologs from *Rattus norvegicus* (rat), *Mus musculus* (mouse), *Schizosaccharomyces pombe* and multiple *Pneumocystis* species (**Supplementary figure 13A)**. We deliberately avoided the C-terminus, which contains the highly conserved histone fold to minimize the risk of cross reactivity with the host histones in partially purified *Pneumocystis* protein preparations. Immunogenic peptides were selected from the N-terminal regional. Peptides with sequence similarities with rat and/or mouse proteins (NCBI accession no. GCF_015227675.2 and GCF_000001635.27, respectively) were excluded using BLASTp (*58*) with word size of 6 and a minimum e-value of 1 as cut off as well as with pattern matches using Perl regular expressions. Additional searches were performed online at https://www.uniprot.org/peptide-search. We select one immunogenic peptide conserved in both *P. carinii* and *P. murina*. Two rabbits were immunized with 20 μg of the affinity purified peptide. The polyclonal antibody was purified by affinity column (GenScript). No cross reactivity against the host cell lysates was seen by western blots (**Supplementary figure 13B**). The following commercial antibodies targeting histones were used at 1:100 dilution for ChIP-Seq: H3 (ab1791, Abcam), H3K4me (39142, Active Motif), H3K9me2 (ab1220, Abcam), H3K9me3 (ab8898, Abcam) H4 (ab1015, Abcam). We verified that peptides used for immunization for commercial antibodies were conserved in *Pneumocystis* proteins.

### Western blots

*Pneumocystis* proteins (antigens) were prepared using glass beads. Partially purified organism pellets were resuspended in 1X PBS buffer. 0.65 ml of 0.5 mm glass beads (Biospec Products, Inc) was added to the suspensions and vortexed for 5 minutes at 4°C. Beads were allowed to settle for less 30 seconds and the supernatant transferred to a new tube for sonication. The samples were centrifuged for 20, 000xg for 10 minutes and the supernatants were transferred to a new tube. The pellet was resuspended in 20 μl PBS.

For western blots, cell lysates and formalin-fixed chromatin preparations were run on Tris-Glycine SDS 4-20% WedgeWell gel (Thermo Fisher Scientific) and transferred on nitrocellulose membranes by incubation at room temperature for 30 minutes, followed by 1 hour incubation in blocking buffer (1X PBS + 5% milk + 0.05% Tween 20) with shaking. Nitrocellulose blots were incubated with primary antibodies (1:100) diluted in blocking buffer for an hour with shaking, followed by a washing and then 1 hour incubation with secondary anti rabbit or anti mouse IgG HRP goat polyclonal antibody (1:2000) (PRF&L) and developed with Pierce 1-Step^TM^ Ultra TMB-Blotting solution (Pierce).

### Immunofluorescence microscopy

For culture, cells were plated in chamber well slide (Millicell EZ slide, Millipore). When cells were ready, the medium was removed and washed with cold PBS 1X, fixed with 4% formaldehyde for 10 minutes at room temperature, and washed three times. Cells were permeabilized with 0.2% Triton X-100 for 30 minutes and followed by a washing with 1X PBST (PBS with 0.05% Tween20).

*Pneumocystis carinii* infected lung tissues fixed in HistoChoice (VWR Life Science) were embedded in paraffin. Five μm sections were labelled with rabbit anti *Pneumocystis* CENP-A antibody (1:200), mouse monoclonal anti *Pneumocystis carinii* antibodies (7C4 and RAE7) (*71, 72*). The 7C4 antibody reacts with one or two uncharacterized antigens of 40 000 daltons and RAE7 reacts with the major surface glycoproteins of *P. carinii* (RAE7 antibody is a gift from Drs. Michael Linke and Peter Walzer, University of Cincinnati College of Medicine) and stained with Vector TrueView autofluorescence quenching kit with DAPI (Vector Laboratories). Alexa Fluor 555 (1:200)– labeled goat anti rabbit IgG (A270039, Invitrogen) and Alexa Fluor 488 (1:200) – labeled goat anti-mouse (A11029, Thermo Fisher Scientific) were used as secondary antibodies. Images were acquired using either a Leica SP8 confocal microscope with x 63x (1.4 NA) HC PL APO objective or a Zeiss 880 confocal microscope with a 63x (1.4 NA) PL APO objective. Images from the Zeiss were taken using 405, 488, and 561 nm excitation with emission bandwidths of 415-480nm, 500-550nm, and 565-700nm. Images from the Leica were taken using 405, 488, and 555 nm excitation with emission bandwidths of 409-491nm, 504-550nm, and 560-721nm. All images were taken with a format of 1024 x 1024 pixels with pixel size ranging from 69-73nm and a line average of 2. Interslice distance for z-stack imaging was set to 160nm and 250nm with pinhole settings of 0.7 airy units and 1 airy unit of images acquired on the Leica and Zeiss microscopes respectively. Image deconvolution was performed assuming an idealized point spread function using Hyugens software program (Scientific Volume Imaging, Netherlands). The software program Imaris (Oxford Instruments, Abingdon UK) was used for image visualization.

*S. pombe* cells were grown overnight at 30°C in minimal medium PMG lacking leucine until logarithmic phase and then mounted in PMG 2% agarose as described (*73*). Images were acquired on a Delta Vision Elite microscope (Applied Precision) with a 100X 1.35 NA oil lens (Olympus). Twenty 0.35-μm z sections were acquired generating maximum intensity projections. Further image processing including background-subtracted centromere foci intensity measurements, was performed using ImageJ (National Institutes of Health).

### ChIP-Seq

Five independent culture experiments with three time points (day 0, 7 and 14) were performed for each species. Purified cells (∼5 x 10^6^ cells) and tissue preparations were fixed by 37% formaldehyde treatment. Chromatin preparation, immunoprecipitation, and Illumina sequencing libraires were prepared using the Low Cell ChIP-Seq kit (Active Motif, Carlsbad, CA) according to the manufacturer instructions. Pre-immune sera served as negative controls (Input). ChIP-Seq libraries were sequenced commercially (Psomagen Rockville, Maryland, USA) using Illumina NovaSeq 6000 (150-bp paired end reads).

### RNA-Seq

Total RNA was extracted using RNeasy Mini kit (Qiagen). RNA quality and integrity were estimated using Bioanalyzer RNA 6000 Pico Assay (Agilent) and sequenced commercially using Illumina HiSeq 4000 (150-base paired end libraries, Novogene Inc, USA). Reads were mapped using STAR v.2.7.9a (*74*), and normalized to bins per million mapped reads (BPM). Transcripts counts were quantified using kallisto 0.44.0 (*75*).

### ChIP-Seq peak calling and sequence analysis

Raw reads were quality-checked with multi-QC (*76*). We removed host DNA contamination from raw sequence reads by mapping against the rat and mouse genomes using Bowtie2 (*77*) with default parameters. Unmapped reads (0.3-5% of total reads) were aligned to the *Pneumocystis* genomes (*P. murina* (GenBank accession number GCA_000349005.2) and *P. carinii* (GCA_001477545.1)), using the Burrows-Wheeler Aligner BWA-MEM (*78*). Alignments were filtered and sorted using SAMTools (*79*) and duplicates marked using PICARD MarkDuplicates http://broadinstitute.github.io/picard). Quality controls were performed using deepTools plotFingerprint (*80*) and phantompeakqualtools (*81*), which estimate the normalized strand coefficient (NSC) and the relative strand correlation (RSC). DeepTools fingerprint plots showed that most reads are mapped within a small fraction of the genome (∼3% of the total 7-8 Mb), which indicates that the antibodies demonstrated a restricted binding (**Supplementary figure 13C)**. ChIP-Seq coverage data were calculated as 50-bp bin and normalized per genomic content (1x normalization). Chromosome ideograms were generated using chromoMap v0.3 (*82*). Enriched regions following immunoprecipitation (IP) were inspected using IGV genome browser.

Because CENP-A is expected to produce sharp peaks CENP-A ChIP-Seq peak calling was performed using MACS3 ChIP enrichment peaks were detected using <u>M</u>odel-Based <u>A</u>nalysis for <u>C</u>hIP-<u>S</u>equencing (*83*) (MACS3 v3.0.0a5) callpeak function with estimated genome sizes and sorted by fold-enrichment (cut off 1.2) and FDR values (broad cutoff 0.1; FDR < 0.05) (raw data are provided in **Supplementary dataset 4**). Enriched peaks were further inspected using deepTools bamCompare versus shuffle non centromeric genomic regions. Peaks were further examined in conjunction with other data (e.g. conservation score, RNA-seq, methylation data, GC content, repeats) with the Integrative <u>G</u>enomics <u>V</u>iewer (*84*), ensuring that peaks are present in all IP samples and absent in controls. Because H3K9me2/3 and H3K4me2 are histone modifications with broad distribution, we used SICER (<u>S</u>patial-clustering method for Identification of <u>C</u>hIP-Enriched <u>R</u>egions) (*85*) with default parameters for peak calling (raw data are provided in **Supplementary dataset 4**).

*De novo* DNA motif searches were performed using MEME v5.1.0 (*86*) and Homer v4.9 (*87*) with a p-value cut off of 10^-10^. GC content across windows (bin) was computed using BEDTools (*88*). Normalized coverage and annotations across chromosomes were visualized using pyGenomeTracks (*89*).

Repeated elements (DNA transposons, retrotransposons and low complexity repeats) were identified using RepeatMasker (*90*) and RepBase (*91*).

Gene located in centromeres were identified by Genes located were extracted from NCBI annotated GFF3 and converted in BED format using custom scripts. Overlaps between gene regions and centromere regions in BED format were identified uysing BEDtools intersect.

### Genome synteny and conservation

DNA alignments were generated using Satsuma2 (*92*) and visualized with Circos (*93*). To estimate the sequence conservation scores, we performed one-to-one pairwise whole genome alignments using LAST (*94*) with the MAM4 seeding scheme (*95*). We consider the trio *P. carinii*/*P. murina*/*P. wakefieldiae.* We generated a neutral model using phyloFit (*96*) from PHAST which fit the tree model to multiple sequence alignments by maximum likelihood using the REV substitution model. Sequence conservation scores were estimated using phastCons (*97*) (--target-coverage 0.25 --expected-length 20 --estimate-trees). Wig files and Bedgraph files were converted using ‘wigToBigWig’ and ‘bigWigToBedGraph’ tools (*98*).

### Bisulfite sequencing

*Pneumocystis* DNA (5 μg) was extracted by a *Pneumocystis* DNA enrichment protocol (*15*). Bisulfite conversion, library preparation and sequencing (Illumina NovaSeq 150 bp paired end reads) were performed commercially (Novogene). Bisulfite conversion of non-methylated DNA was performed using EZ DNA Methylation kit. Data analysis was performed using BSMAP and methratio script (*99*). Only cytosine positions with more than 5-fold coverage were considered. We used weighted methylation percentage (*44*), which is calculated as the number of reads supporting methylation over the number of cytosines sequenced to quantify methylation levels. Methylation data were partitioned over different genomic compartments using BEDTools (*100*). Statistical comparisons were performed using R (https://www.R-project.org) to compute non-parametric Mann-Whitney U test and Bonferroni correction to adjust *P*-values for multiple comparisons.

### ChIP-qPCR

CENP-A enrichment and H3 depletion relative to H4 within the centromeres were evaluated using primers specific to centromeric and non-centromeric loci (control region taken 200 kilobases away from centromere 1 of each species) (see **Supplementary Table 5** for primers). Primers were designed using primer3 (*101*) from regions visualized using IGV genome browser. Primers with potential matches against mammal genomes were excluded using NCBI BLASTn. Dilutions of 1:10 were used based on preliminary test runs. Each reaction contains 5 μl of iTaq universal Sybr green supermix (Biorad), 1.25 μl of primers (500 nM)) and 2.5 μl of DNA. All assays were run on CFX96 thermocycler (Biorad). Three technical replicates were taken for each assay, and the standard errors of the mean were calculated. The PCR program was as follow: initial denaturation for 2 min at 95°C, followed by 30 cycles of 30 seconds at 95°C, 60°C for 30s, and 72° for 30s. Locus ΔCt values were normalized using the formula: (^ΔΔCt^ ChIP – ^ΔCt^ Input; https://www.sigmaaldrich.com/US/en/technical-documents/protocol/genomics/qpcr/chip-qpcr-data-analysis) and the fold enrichment of the ChIP DNA over the input was computed as log2 (2^ΔΔCt^). The plots and statistical analyses were performed with GraphPad Prism 8.

### ChIP-LC-MS

Formalin-fixed purified chromatin preparations were separately co-immunoprecipitated with anti CENP-A, H4 and pre immune serum (control) using Pierce™ Direct Magnetic IP/Co-IP Kit (ThermoFisher). Each IP reaction was performed separately on aliquots from the same chromatin preparation. Elution was performed using 1% formic acid. Extracts were frozen in dry ice, lyophilized for 1 hour and digested using trypsin, diluted, and injected to a Thermo Orbitrap Fusion LC-MS/MS to identify unique peptides. LC-MS/MS data were searched against the proteomes using Proteome Discoverer 2.4 (Thermo Fisher Scientific, Waltham, MA). Label-free quantification was analyzed based on the peak intensity of the precursor ion.

## Acknowledgments

We thank Rene Costello for animal studies, Drs. Keytam Awad, Salina Gairhe and Shuibang Wang for discussion about ChIP-Seq. This work has been funded in whole or in part with federal funds from the Intramural Research Program of the US National Institutes of Health (NIH) Clinical Center, the National Institute of Allergy, and Infectious Diseases (NIAID) and the National Cancer Institute. The views expressed in this work are those of the authors and do not necessarily represent the official positions of the NIH or U.S. Government. This study used the Office of Cyber Infrastructure and Computational Biology (OCICB) High Performance Computing (HPC) cluster at the National Institute of Allergy and Infectious Diseases (NIAID), Bethesda, MD. This study also utilized the high-performance computational capabilities of the Biowulf Linux cluster at the NIH, Bethesda, MD (http://biowulf.nih.gov).

## Author contributions

Conceptualization: OHC, JAK

Methodology: OHC, HDF, JAK, SC

Investigation: OHC, SC, HDF, YL, LB, HW, ERF. ASD, ST, JPD, MC, LM

Visualization: OHC, JAK, SC, LB, CB, HDF

Supervision: OHC, JAK, SG

Writing—original draft: OHC

Writing—review & editing: OHC, JAK, HDF, LM, SC, YL, LB, HW, ERF, ASD, CB, ST, JPD, SG, MC

## Data and materials availability

ChIP-Seq datasets and associated peak calling outputs, RNA-seq and bisulfite sequencing data generated in this study are deposited at NCBI GEO (accession numbers GSE232453, GSE226536, GSE232357, GSE232355, GSE230598). Codes generated for this project are available at GitHub: https://github.com/ocisse/pneumo-cens

## Supplementary Materials

Figs. S1 to S10

Figs. S11 to S13 and Tables S1 to S4 are available at DOI: https://doi.org/10.5281/zenodo.10574230

